# Utility of an untargeted metabolomics approach using a 2D GC-GC-MS platform to distinguish relapsing and progressive multiple sclerosis

**DOI:** 10.1101/2024.02.07.579252

**Authors:** Indrani Datta, Insha Zahoor, Nasar Ata, Faraz Rashid, Mirela Cerghet, Ramandeep Rattan, Laila M Poisson, Shailendra Giri

**Affiliations:** Department of Public Health Sciences, Henry Ford Health, Detroit, MI, 48202, USA; Department of Neurology, Henry Ford Health, Detroit, MI, 48202, USA; Women’s Health Services, Henry Ford Health, Detroit, MI, 48202, USA; Department of Neurosurgery, Henry Ford Health, Detroit, MI, 48202, USA

**Keywords:** GC-GC-MS, metabolomics, multiple sclerosis, RRMS, PPMS

## Abstract

**Introduction:** Multiple sclerosis (MS) is the most common inflammatory neurodegenerative disease of the central nervous system (CNS) in young adults and results in progressive neurological defects. The relapsing-remitting phenotype (RRMS) is the most common disease course in MS and may progress to the progressive form (PPMS).

**Objectives:** There is a gap in knowledge regarding whether the relapsing form can be distinguished from the progressive course or healthy subjects (HS) based on an altered serum metabolite profile. In this study, we performed global untargeted metabolomics with the 2D GCxGC-MS platform to identify altered metabolites between RRMS, PPMS, and HS.

**Methods:** We profiled 235 metabolites in the serum of patients with RRMS (n=41), PPMS (n=31), and HS (n=91). A comparison of RRMS and HS patients revealed 22 significantly altered metabolites at p<0.05 (false discovery rate [FDR]=0.3). The PPMS and HS comparisons revealed 28 altered metabolites at p<0.05 (FDR=0.2).

**Results:** Pathway analysis using MetaboAnalyst revealed enrichment of four metabolic pathways in both RRMS and PPMS (hypergeometric test p<0.05): 1) galactose metabolism; 2) amino sugar and nucleotide sugar metabolism; 3) phenylalanine, tyrosine, and tryptophan biosynthesis; and 4) aminoacyl-tRNA biosynthesis. The Qiagen IPA enrichment test identified the sulfatase 2 (SULF2) (p=0.0033) and integrin subunit beta 1 binding protein 1 (ITGB1BP1) (p=0.0067) genes as upstream regulators of altered metabolites in the RRMS vs. HS groups. However, in the PPMS vs. HS comparison, valine was enriched in the neurodegeneration of brain cells (p=0.05), and heptadecanoic acid, alpha-ketoisocaproic acid, and glycerol participated in inflammation in the CNS (p=0.03).

**Conclusion:** Overall, our study suggested that RRMS and PPMS may contribute metabolic fingerprints in the form of unique altered metabolites for discriminating MS disease from HS, with the potential for constructing a metabolite panel for progressive autoimmune diseases such as MS.

## Introduction

Multiple sclerosis (MS) is an autoimmune disorder of the central nervous system (CNS) that is characterized mainly by immune cell infiltration, inflammation, and demyelination (Lucchinetti et al., 2000). The disease can occur in younger adults between the ages of 20 and 50 years. Relapsing-remitting MS (RRMS) is the most common type of MS and comprises ∼85% of the diagnoses (Oh et al., 2008). MS relapse was defined as a relapse of MS with periods of remission occurring in between. Approximately 90% of RRMS patients eventually progress to secondary-progressive MS (SPMS) (Garg et al., 2015; Dobson and Giovannonic, 2019). Approximately 10-15% of patients diagnosed with MS progress from the beginning, termed primary progressive MS (PPMS), and their neurological function declines much faster than that of any other MS type. Approximately half of individuals with a mild disease phenotype progress to a secondary progressive phenotype within 10 years. Overall, MS diagnosis encompasses the integration of clinical, imaging, and laboratory findings, as there is still no single reliable clinical feature or diagnostic laboratory biomarker. Furthermore, there is no single blood-based diagnostic test that can diagnose MS and discriminate between RRMS and PPMS (Harris et al., 2017). Identification and confirmation of such blood-based diagnostic tests will be highly beneficial for diagnosing this disease and its progression.

With the advancement of high-throughput molecular omics platforms, metabolomics has emerged as a highly beneficial technology with tremendous potential for the detection of therapeutic strategies for MS (Zahoor et al., 2021). Metabolomics is the study of the metabolome within cells, biofluids, or tissues to identify and quantify small-molecular-weight metabolites. Using biostatistical and bioinformatics tools, metabolomics allows the identification of metabolic pathways that could be targeted for the development of therapies. Several studies have employed targeted and untargeted metabolomics using MS biofluids to identify metabolic changes during the disease course (Zahoor et al., 2021). In this study, we used two-dimensional GCxGC/MS, which can detect a much greater number of chromatographic peaks at a lower detection limit for small molecules in various biological mixtures than GC/MS or NMR (Storey et al., 2020). Here, we provide a comprehensive untargeted metabolomics analysis of RRMS and PPMS patients via the 2D GCxGC/MS platform in comparison to the respective control subjects. In addition to commonly altered metabolites, significantly altered associated pathways may explain the difference in disease course between RRMS and PPMS patients. Ultimately, such a metabolite panel would support clinical care after appropriate validation in large-scale samples. This approach could lead to the generation of a specific metabolite signature for differentiating between relapsing and progressive MS phenotypes. This study has immense potential to aid in the early detection of MS in the clinical setting and could aid in identifying the disease in its early stages and preventing its progression to a more severe course accompanied by disability, thereby rescuing the young, productive population from becoming crippled. This would eventually mean saving millions of dollars in investment in the U.S. healthcare system and lessening the economic burden of the disease.

## Materials and methods

### Human subjects

Deidentified serum samples were obtained from the repository of the Accelerated Cure Project (ACP). All basic demographic information (age, sex, race, and ethnicity) and diagnostic groups (RRMS, PPMS, and HS) were collected from the medical records of the ACP. The human participants were recruited for this study by ACP after written informed consent was acquired from them following the ethical standards established by the World Health Organization (WHO) and the Declaration of Helsinki 1964 and its later amendments or comparable ethical standards. Serum samples were acquired from ACP through a Henry Ford Health-approved IRB study of the metabolomics signature in MS patients. The sample numbers for the RRMS group were 41, 31 for PPMS patients, and 91 for HS. The demographic details of the study subjects are given in Table 1.

**Table 1:** Demographics and characteristics of relapsing-remitting (RRMS) patients, progressive MS (PPMS) patients, and matched healthy individuals (HS)

| Variable |  | RRMS | CTRL for RRMS | PPMS | CTRL for PPMS |
| --- | --- | --- | --- | --- | --- |
| N |  | 41 | 44 | 31 | 47 |
| Age, median [IQR] |  | 39 [34, 48] | 39.5 [33.75, 49] | 49 [46, 58.5] | 53 [46.5, 60.5] |
| Sex, N (%) | Male | 12 (29.3) | 13 (29.5) | 9 (29.0) | 12 (25.5) |
|  | Female | 29 (70.7) | 31 (70.5) | 22 (71.0) | 35 (74.5) |
| Race<sup>+</sup>, N (%) | Black | 4 (9.8) | 4 (9.1) | 1 (3.3) | 1 (2.1) |
|  | White | 35 (85.4) | 39 (88.6) | 28 (90.3) | 44 (93.6) |
|  | Other<sup>*</sup> | 2 (4.8) | 1 (2.3) | 2 (6.4) | 2 (4.3) |

\* Other includes “American Indian or Alaskan Native”, “Asian, other than Middle East or South Asian”, and “Native Hawaiian or Pacific Islander.” The categories are collapsed here to protect patient identities.
+ No patients were identified as “Hispanic or Latino”, so ethnicity was not separately reported.

### Sample preparation

All samples were processed in random order and were blinded to the sample group to avoid systemic bias. Then, 400 µL of 100% methanol solution was added to 100 µL of sample. The mixture was vortexed for 2 min and then placed on ice for 15 min. The sample was then centrifuged at 4°C and 15000 rpm for 20 min. After 300 µL of the supernatant was transferred into a glass vial, the transferred supernatant was first dried by a Speedvac to remove methanol, followed by freeze-drying to remove water. Each metabolite extract was then dissolved in 30 µL of pyridine with 20 mg/mL methoxyamine hydrochloride and vigorously vortexed for 1 min. Methoxymation was carried out by sonicating the solution for 20 min followed by 1 h of incubation at 60°C. Derivatization was conducted by adding 30 µL of N-methyl-N-(trimethylsilyl)trifluoroacetamide (MSTFA). The solution was incubated at 60°C for another 1 h. The stock solutions were then transferred to GC vials for analysis. The methoxymation and derivatization were carried out just before GC×GC-TOF MS analysis. Three pooled samples were prepared simultaneously by mixing 50-100 µL sample supernatants and then conducting methoxymation and derivatization. Pooled samples were analyzed via GC×GC-TOF MS after analysis of every five biological samples.

### GC×GC-TOF MS analysis

A LECO Pegasus GC×GC-TOF MS instrument was coupled with an Agilent 6890 gas chromatograph and a Gerstel MPS2 autosampler (GERSTEL, Inc., Linthicum, MD), featuring a LECO two-stage cryogenic modulator and secondary oven. The primary column was a 60 m × 0.25 mm ^1^d_c_ × 0.25 µm ^1^d_p_ DB-5 ms GC capillary column (a phenyl arylene polymer virtually equivalent to 5%-phenyl-methylpolysiloxane). The secondary GC column (1 m × 0.25 mm ^1^d_c_ × 0.25 µm ^1^d_f_, DB-17 ms (50% phenyl)-methylpolysiloxane) was placed inside the secondary GC oven following the thermal modulator. Both columns were obtained from Agilent Technologies (Agilent Technologies J&W, Santa Clara, CA). The helium carrier gas (99.999% purity) flow rate was set to 1.0 mL/min at a corrected constant flow via pressure ramps. The inlet temperature was set at 280°C. The primary column temperature was programmed at an initial temperature of 60°C for 0.5 min, ramped at 5°C/min to 270°C, and maintained for 11 min. The secondary column temperature program was set to an initial temperature of 70°C for 0.5 min and then ramped at the same temperature gradient employed in the first column to 280°C. The temperature of the thermal modulator was set to +15°C relative to the temperature of the primary oven, and a modulation time of P_m_ = 2 s was used. The mass range was set to 29-800 *m/z*, and the acquisition rate was 200 mass spectra/second. The ion source chamber was set at 230°C with a transfer line temperature of 280°C, and the detector voltage was set at 1390 V with an electron energy of 70 eV. The acceleration voltage was turned on after a solvent delay of 544 seconds. The split ratio was set at 10:1.

### Data extraction and compound identification

LECO’s instrument control software ChromaTOF was used to process the GC×GC-TOF MS data for peak picking and tentative metabolite identification, followed by retention index matching, peak merging, peak list alignment, and normalization. For metabolite identification using ChromaTOF, each chromatographic peak was tentatively assigned to a metabolite if its experimental mass spectrum and a database spectrum had a spectral similarity score of no less than 500 (a maximum spectral similarity score: 1000). Peak merging and peak list alignment were carried out using MetPP software, while retention index matching was performed using iMatch with the *p-value* set as *p* ≤ 0.001.

### Statistical analysis

Missing intensity values, indicating a technical error or low metabolite levels among sample groups, were imputed with the KNN algorithm. This step was followed by normalization (Johnson transformation) and batch correction (ComBat, Johnson et al., 2007) of the metabolite intensities. Principal component analysis was performed to analyze the remaining samples. Partial least squares discriminant analysis (PLS-DA) was used for the assessment of the separability of the samples (**Figure 1**). T-tests, allowing unequal variance, were used to compare changes in mean expression, per metabolite, between patients with RRMS or PPMS and the corresponding healthy control group (HS). P values <0.05 were considered to indicate statistical significance and were visualized with a heatmap. Due to multiple testing t-tests, p values were transformed into q values (Storey et al., 2020), both of which are reported. KEGG pathway analysis (http://www.genome.jp/kegg) of 80 *Homo sapiens*-associated pathways considered both the statistical enrichment of changed intensity using GlobalTest (Xia and Wishart, 2016) and the impact of metabolite changes based on pathway topology using the relative betweenness centrality measure (Xia and Wishart, 2016). Further pathway enrichment testing, based on Fisher’s exact tests, and the construction of metabolite networks were conducted using Qiagen’s IPA knowledgebase (Bento et al., 2011). To build a classifier, multivariate feature selection was conducted with the biosigner method (Rinaudo et al., 2016), which uses SVM, PLS-DA, and random forest classifiers in parallel to create a signature for binary classification. In brief, the dataset is partitioned into training and testing sets. Each of these models is trained on a training set, and prediction accuracy is evaluated in a testing set, with metabolites ranked based on importance. Finally, the biosigner method returns a tier for each feature (metabolite) per classifier, with tier S indicating inclusion in the final signature after all steps and other tiers (A to E) indicating less preference. Except where noted, statistical analyses were conducted with “R” (http://cran-r-project.org/) or “MetaboAnalyst 5.0” (http://www.metaboanalyst.ca) (Xia and Wishart, 2016).

**Figure 1:**
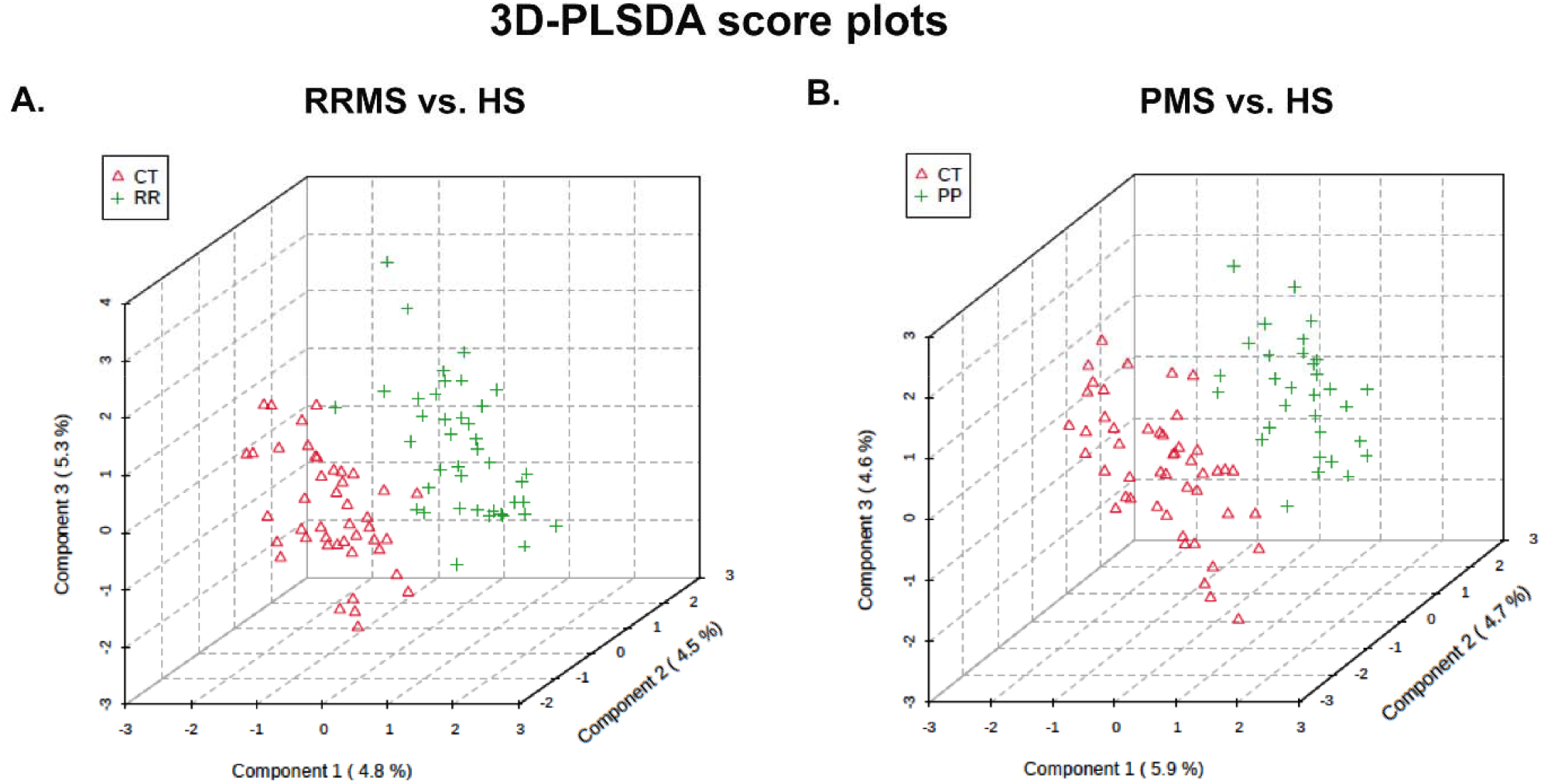
Partial least squares discriminant analysis (PLS-DA) PLSDA plot showing the results between (A) RRMS and HS and (B) PPMS and HS for the X and Y matrices.

## Results

The demographic data in **Table 1** show that the median age of the RRMS patients was 39 years, whereas the median age of the PPMS patients was 49 years. Twenty-nine patients were females in the RRMS group, and 22 patients were females in the PPMS group, with an overall preponderance of MS in females (70.7-71%). Most of the MS patients were white, with 85.4% having RRMS and 90.3% having PPMS. The global metabolomic profiles of RRMS, PPMS, and HS serum samples were generated via fine mapping via a 2D GC□GC□MS platform. A total of 235 structurally different biochemicals were detected among these samples (**Table S1, S2, S3**). PLS-DA showed a clear separation of the metabolites between RRMS and HS and between PPMS and HS (**Figure 1**). Between the RRMS patients and controls, 20 metabolites were significantly altered (8.5% of the 235 metabolites detected), with 11 metabolites increasing and 9 metabolites decreasing in the RRMS patients relative to the controls (p<0.05, with an FDR of 0.3) (**Table S1)**. Between the PPMS patients and controls, 26 metabolites were significantly altered (10.6% of the 235 metabolites detected), with 10 metabolites increasing and 16 metabolites decreasing in the PPMS patients relative to the controls (p<0.05, with an FDR of 0.2) (**Figure 2, Table S3**). To visualize the relationships between the altered metabolites, heatmaps were drawn using hierarchical clustering (**Figure 2A, 2B**). Six metabolites were common between these two comparisons (**Figure 2C, Table S3**), and the changes were directionally consistent for RRMS and PPMS relative to their respective control groups (**Figure 2D**). These common metabolites included methyl 11,14-eicosadienoate (S), 11,14-eicosadienoic acid, L-tyrosine, 2-hydroxypentanoic acid (S), erythrose, and margaric acid (C17) (**Figure 2D, Table S3**).

**Figure 2:**
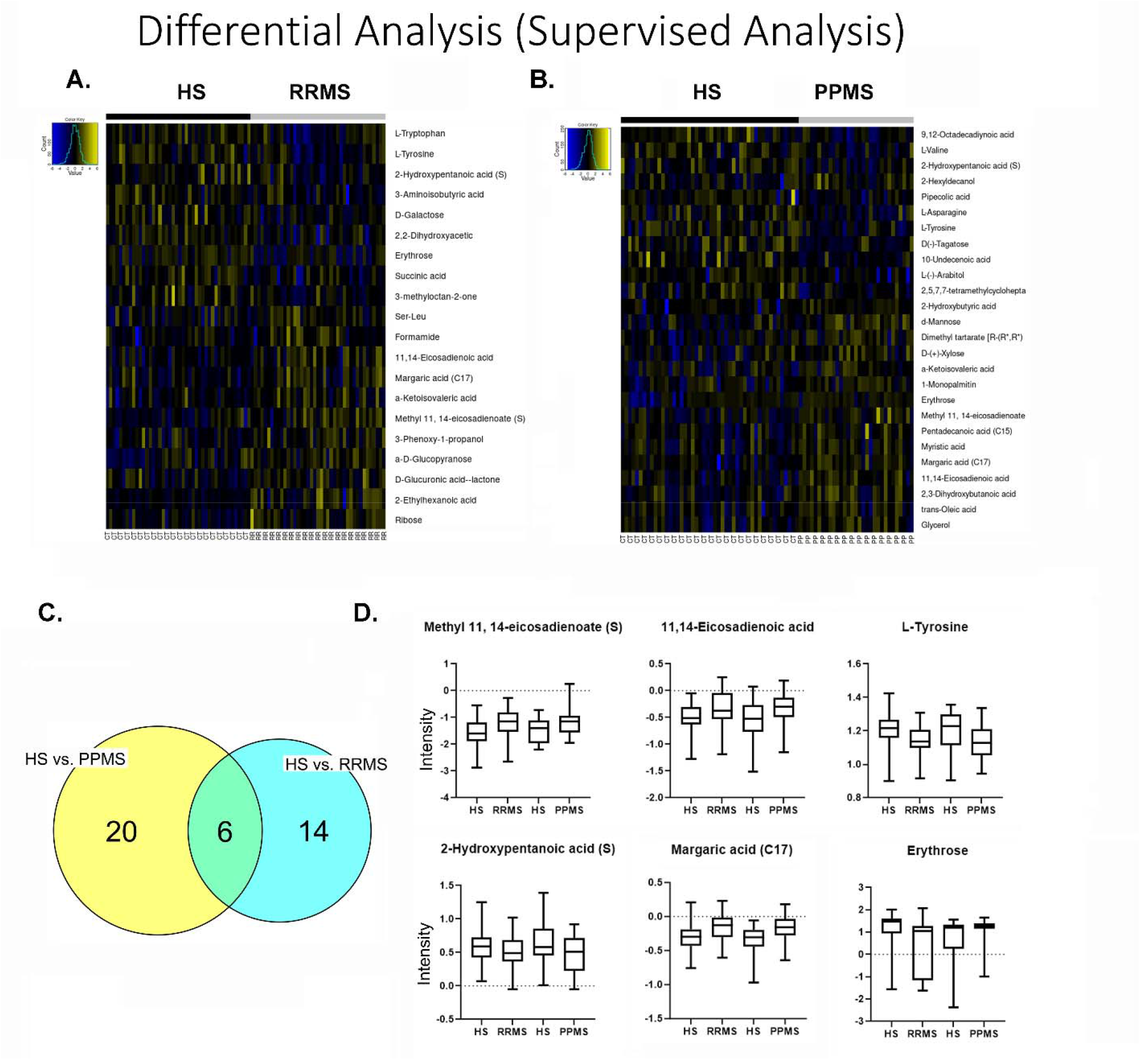
Metabolite profile of metabolites across HS, RRMS, and PPMS. (A) Heatmap of differential metabolites between RRMS patients and HS. (B) Heatmap of differential metabolites between PPMS and HS. (C) Overlap between differential metabolites between RRMS and PPMS patients. (D) Intensity plots depicting the differences among the HS, RRMS, and PPMS patients.

To understand the functional role that these altered metabolites may play in the serum, the KEGG metabolic library was analyzed using MetaboAnalyst (Chong et al., 2019; Xia and Wishart, 2016). The results of each of the 80 human pathways of KEGG were simultaneously tested to determine the most significant pathways in terms of hypergeometric test p values <0.05. The top four pathways according to the p-value (top four) were identified as follows: 1) aminoacyl-tRNA biosynthesis; 2) phenylalanine, tyrosine, and tryptophan biosynthesis; 3) amino sugar and nucleotide sugar metabolism; and 4) galactose metabolism (**Figure 3A**). In the PPMS and control groups, the top four pathways included 1) phenylalanine, tyrosine, and tryptophan biosynthesis; 2) galactose metabolism; 3) pentose and glucuronate; and 4) aminoacyl-tRNA biosynthesis, with the most significant pathways identified in terms of hypergeometric test p-values <0.05 being similar to those mentioned for the RRMS and control groups, but aminoacyl-tRNA biosynthesis was more significant (**Figure 3B**). An enrichment test with the Qiagen IPA knowledgebase identified sulfatase 2 (SULF2) (low in RRMS) and integrin subunit beta 1 binding protein 1 (ITGB1BP1) (high in RRMS) as upstream regulators at p values of 0.0033 and 0.0067, respectively, with an activation z score of 2 (absolute value) in RRMS patients and controls (**Figure 3C**). However, in the PPMS and control groups, valine was enriched in the neurodegeneration of brain cells at a p-value of 0.05, and heptadecanoic acid, alpha-ketoisocaproic acid, and glycerol participated in inflammation in the CNS at a p-value of 0.03 (**Figure 3D**).

**Figure 3:**
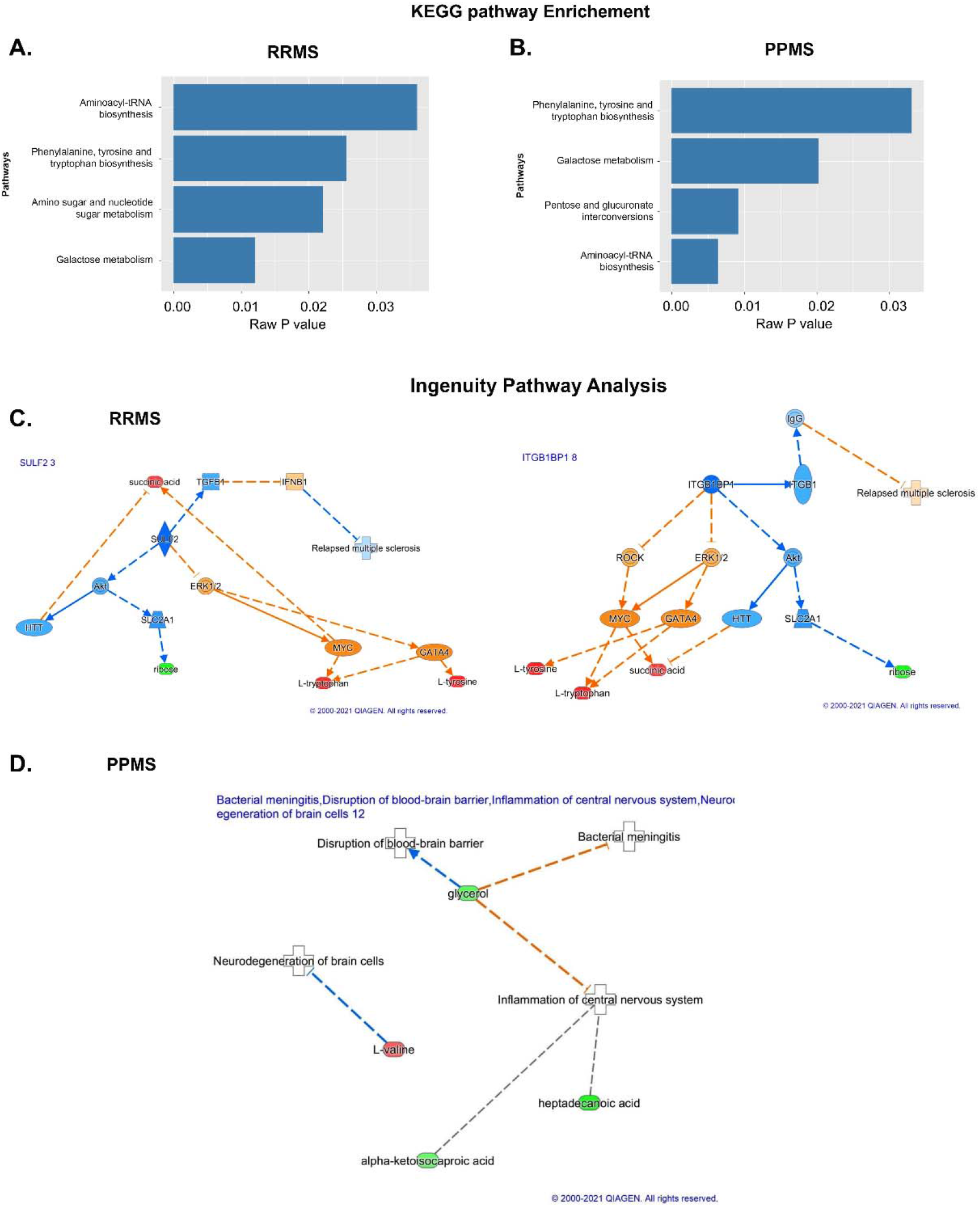
Pathway analysis using KEGG and IPA. (A&B) KEGG pathway analysis in which the pathways and raw p values are shown for the RRMS and PPMS patients. (C&D) IPA of the molecules SULF2 3 and ITGB1BP1 8 in RRMS and PPMS. Associations of differentially expressed metabolites between RRMS and HS with biological processes and upstream regulators determined via IPA. In addition, SULF2 3 and ITGB1BP1 8, which are the primary regulators activated for activation based on the different levels of metabolites in the chain below, exhibited connectivity. The prediction of activation is shown by the broken orange lines, whereas the prediction inhibition is represented by the broken blue line. The red color shows the upregulation of RRMS. The last layer shows the different metabolites in RRMS and HS.

The PLS-DA algorithm that we utilized is flexible and can be applied to both descriptive and predictive modeling. On the other hand, the support vector machine (SVM) is a supervised learning model that works with associated learning algorithms to analyze data for classification. Finally, the random forest algorithm integrates the outputs from several decision trees to produce a single result while minimizing the opportunity for overfitting. Among the three models we tested in this class (RRMS vs HS) and RRMS-the SVM was the most accurate, focused on the results, with an accuracy rate of 77%. Multivariate machine learning with PLS-DA, random forest, and SVM identified 12 metabolites that can differentiate between RRMS patients and controls at 73%, 75%, and 77%, respectively (**Figure 4A, Table 2**), and the levels or intensities of these metabolites in RRMS patients are presented as a bar graph (**Figure 4B**). These 12 metabolites included 2-ethylhexanoic acid, ribose, erythrose, 3-indole-propionic acid, a-D-glucopyranose, D-glucuronic acid-lactone, heptanoic acid (C7), L-threose (syn), lanthionine, linoleic acid 9,12-octadecadienoic acid, succinic acid, and methyl 11,14-eicosadienoate. The overall RRMS ‘S’-signature status prediction was low for the PPMS (**Figure 5**).

**Table 2:** The accuracy of different models for control and RRMS patients.

| Name of Classifier | PLS-DA | Random Forest | SVM |
| --- | --- | --- | --- |
| Accuracy | 73% | 75% | 77% |

**Figure 4:**
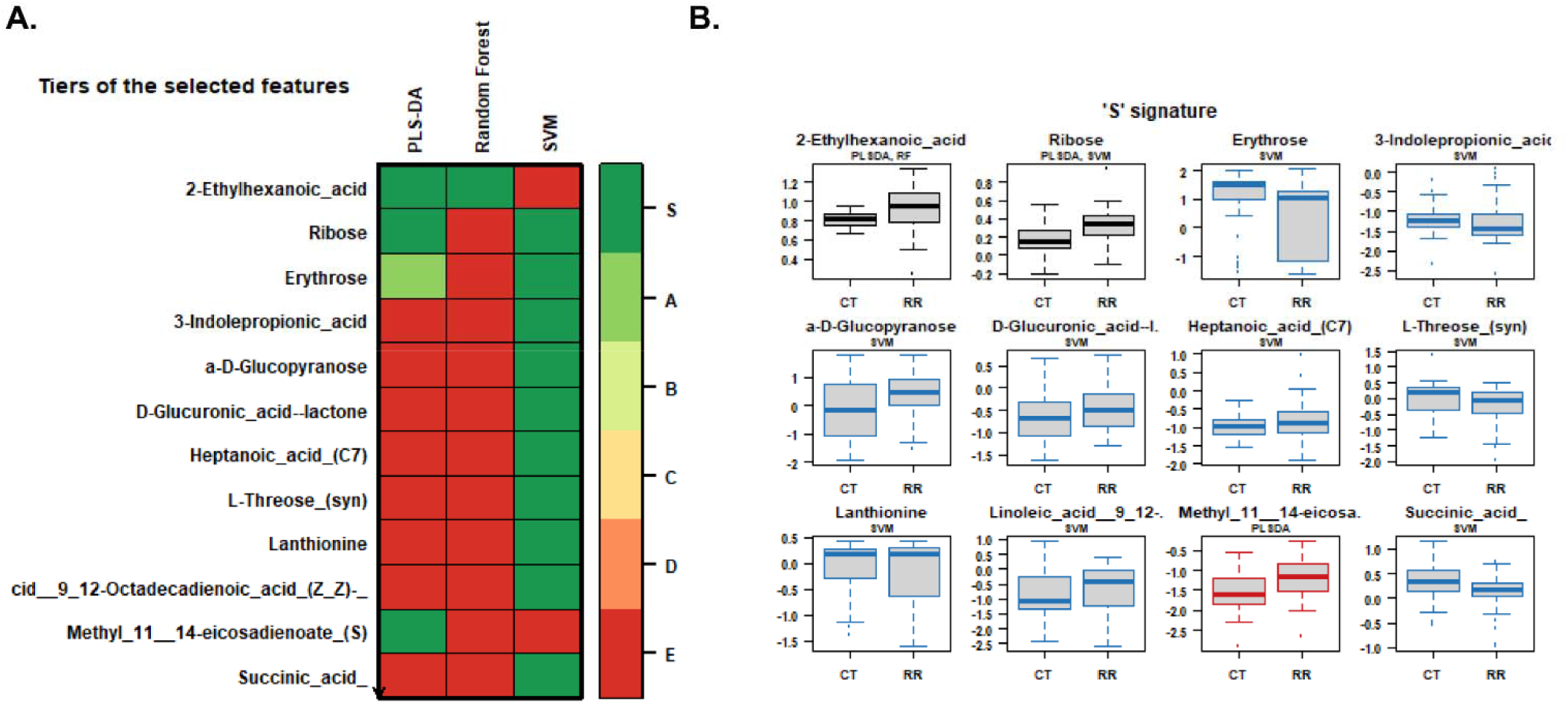
Heatmap and signature of metabolites. (A) Different tiers of selected features with PLS-DA and SVM random forest to binary classifiers on a scale of red to green showing their intensities to select a smaller subset of metabolites with the highest predictive accuracies. (B) S signatures of different metabolites identified through the classifiers for control and disease patients.

**Figure 5:**
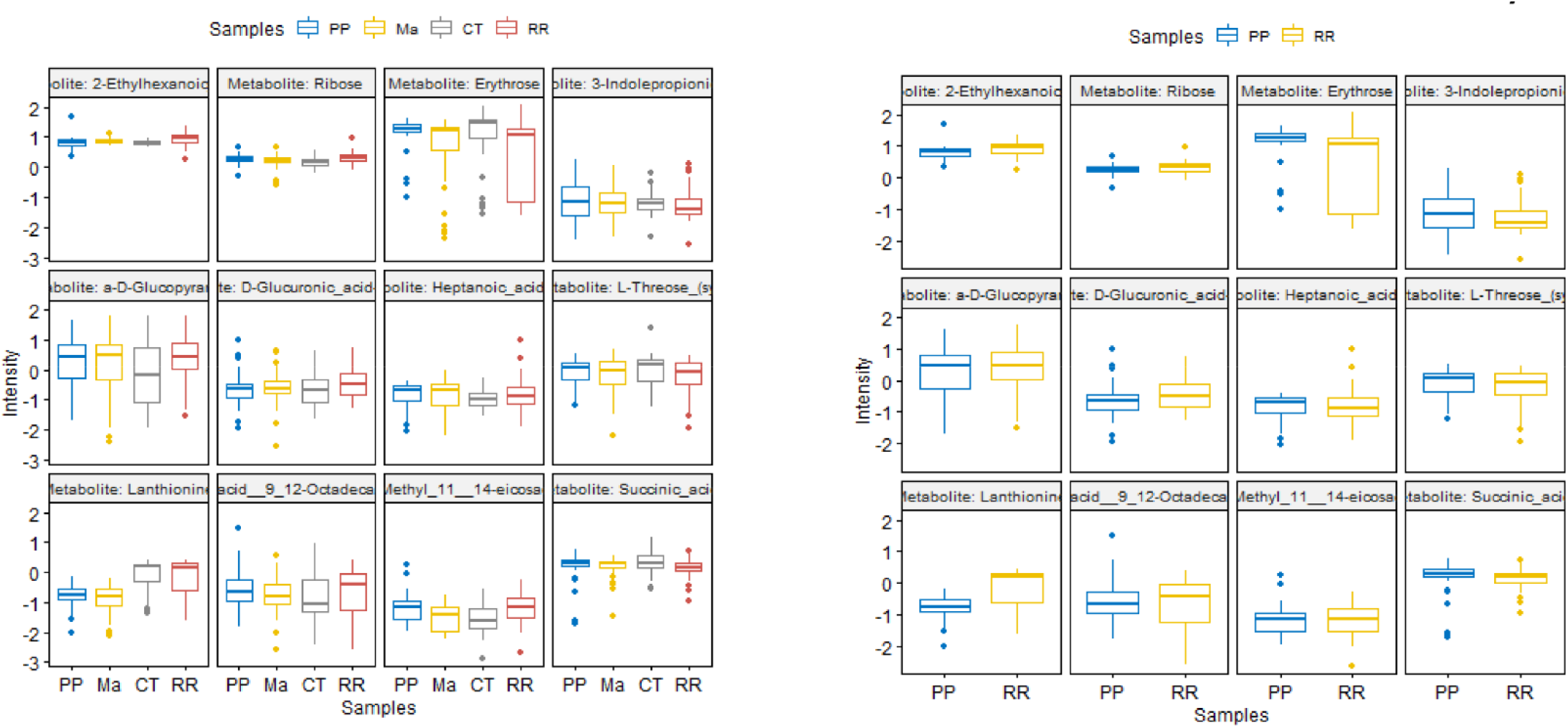
S-signature across the HS, RRMS, and PPMS cohorts. The RRMS signature obtained by binary classifier intensities in PPMS patients and healthy controls, prediction of the metabolites, and prediction via the PPMS were low.

## Discussion

Treatment of MS in the early stage, such as RRMS, is more beneficial for delaying disability than treatment in the more advanced progressive stage (Harris et al., 2017). Several drugs have been approved for the treatment of MS, but most of their ability to reduce the onset of disability is still under evaluation (Garg et al., 2015; Dobson and Giovannonic, 2019). In addition, there are several efforts underway to search for biomarkers in biofluids with different high-dimensional omics platforms in MS. The main aim of this study was to investigate altered metabolites in different pathways corresponding to the RRMS and PPMS stages of MS with an advanced global GC-GC□MS metabolomics platform. Disease mechanisms can be discerned via alterations in pathways by comparing RRMS patients with PPMS patients. KEGG and enrichment pathway analysis of the RRMS patients and controls revealed galactose metabolism and amino sugar and nucleotide sugar metabolism, with alpha-D-glucose and alpha-D-galactose as significant metabolites altered in these pathways. The level of alpha D-galactose was lower in the RRMS group than in the HS group. It is a monosaccharide and an essential component in galactose metabolism and enters glycolysis through its conversion to glucose-1-phosphate by multiple steps from the Leloir pathway. This glucose-1-phosphate is formed by the protein galactokinase encoded by the GALK1 gene. Interestingly, other studies have shown that the administration of galactose improved the performance of patients by promoting remyelination. This type of galactose administration is beneficial for reducing other neurogenerative diseases, such as Alzheimer’s disease (Ravera and Panfoli, 2015). Importantly, in our study, we observed a reduction in the alpha-D-galactose concentration, suggesting degradation of the myelin sheath in RRMS patients compared to healthy subjects. Alpha-D-glucose, another altered metabolite that participates in these pathways, is a hexose organic compound and a monosaccharide. We observed that alpha-D-glucose was elevated in RRMS patients compared to HS. Elevated glucose levels could be an indication of insulin resistance, and many peripheral insulin resistance diseases can impair brain structure and function and lead to cognitive impairment (de la Monte, 2017). Our finding of elevated D-glucose could subsequently suggest cognitive impairment in RRMS patients. The other two significantly affected pathways were phenylalanine, tyrosine, tryptophan biosynthesis, and aminoacyl-tRNA biosynthesis, which were enriched with L-tyrosine and L-tryptophan as the altered metabolites in these pathways.

Tyrosine is a nonessential amino acid that is generated from L-phenylananine by phenylalanine hydroxylase and metabolized into catecholamine neurotransmitters (Chandel, 2021). We observed a lower level of L-tyrosine in RRMS in our study, suggesting a reduction in the biosynthesis of these neurotransmitters. L-tyrosine has also been associated with metabolic syndrome and could be an early biomarker for this disease (Hellmuth et al., 2016). Similarly, the L-tryptophan concentration was lower in the RRMS group than in the HS group. Likewise, L-tryptophan is an essential protein amino acid that bears an indole ring, and its derivatives lead to the synthesis of the neurotransmitter hormone serotonin (5-HT), penial grand melatonin, and the trace amine tryptamine (Chandel, 2021). Abnormalities in 5-HT synthesis are related to the pathophysiology of many neurological disorders, such as mood disorders, Parkinson’s disease, sleep disorders, dementia, Huntington’s disease, and Tourette’s syndrome. This amino acid also participates in the kynurenine pathway. This pathway is involved in the synthesis of nicotinamide adenine dinucleotide (NAD) and is upregulated by neurogenerative triggers (Palego et al., 2016; Blankfield, 2012; Sandyk, 1992). These findings suggest that alteration of L-tryptophan leads to CNS dysfunction. Thus, the phenylalanine, tyrosine, and amino sugar pathways play significant roles in the pathology of MS.

In the PPMS cohort, aminoacyl-tRNA biosynthesis was the most significantly altered pathway, and L-asparagine, L-valine, and L-tyrosine were altered in this pathway. The serum levels of all three of these metabolites were elevated in the PPMS group compared to the HS group. L-asparagine is a nonessential common amino acid that contains side chain carboxamide, and oxaloacetate is the precursor. L-Asparagine is already known for its use in chemotherapeutic applications, but its neuroprotective effect on Parkinson’s disease has been shown in a study in which it was used as a cell model because it activates autophagy and mitochondrial fusion (Zhang et al., 2020). L-asparagine is also known for its immunosuppressive and anti-inflammatory properties. L-valine is an essential proteinogenic branch chain amino acid and is associated with maple syrup-related urine disease (Manoli and Venditti, 2016). In sickle-cell disease, a single glutamic acid (hydrophilic) in beta-globin is exchanged with valine (hydrophobic), resulting in abnormal hemoglobin aggregation (https://themedicalbiochemistrypage.org/galactose-metabolism/).

Furthermore, 6 significantly altered metabolites overlapped between the RRMS and PPMS patients: methyl 11,14-eicosadienoate (S), L-tyrosine, 11,14-eicosadienoic acid, margaric acid (C17), erythrose, and 2-hydroxypentanoic acid (S). Most of these altered metabolites belong to the polyunsaturated fatty acid (PUFA) group. A study of human brains from postmortem patients with moderate and severe Alzheimer’s disease and dementia with Lewy bodies (DLB) identified 24 fatty acids (Nasaruddin et al., 2018). Among those identified, fatty acids, such as 11,14-eicosadienoic acid, were present at higher levels in patients with moderate Alzheimer’s disease and DLB than in patients with severe Alzheimer’s disease. This finding suggested a relationship between lipid metabolism and disease pathology. A higher brain fatty acid content also leads to ceramide accumulation, which can increase amyloid beta peptide levels. Similarly, in our study, we observed that 11,14-eicosadienoic acid was more abundant in RRMS patients than in PPMS patients. In addition, another study showed that eicosadienoic acid can alter the response of macrophages to inflammatory stimulation (Huang et al., 2011). Likewise, many studies have shown that PUFAs are highly enriched in the CNS. The benefits of alcohol intake include many psychiatric and neurological disorders, including neurodegenerative conditions (Dyall and Michael-Titus, 2008). Moreover, the reduced level of ethyrose in RRMS patients observed in comparison to that in PPMS patients in our study indicates that in many other neurogenerative diseases, such as Alzheimer’s disease, a reduction in glucose metabolism leads to disease progression. This finding suggested that the decreased intensity of erythrose in RRMS patients may intensify their progression to more advanced stages of MS and could lead to disease progression (Gibson et al., 2013).

Taken together, the results of the present study explored the alteration of metabolites in RRMS and PPMS using an advanced 2D GC-GC□MS platform via significantly affected pathways for the development of a targeted metabolite panel to monitor disease progression in MS. The specific altered metabolites and associated pathways found in MS in the present study reflect their role in myelination, maintenance of brain structure, cognitive function, and inflammation, as confirmed by reports on other neurodegenerative disorders. However, further validation of these altered metabolites in a large sample cohort is needed before any clinical use of these materials.

## Author Contributions

ID performed the analysis and wrote the first draft. IZ refined the manuscript and compiled the final version. LMP and NA provided inputs in the data analysis. LMP, MC, FR, and RR edited the manuscript. SG directed the study, reviewed the data analysis, and edited the manuscript before approval was obtained for submission.

## Compliance with Ethical Standards

### Ethics Approval

The human participants were recruited for this study after written informed consent was acquired from them following the ethical standards established by the World Health Organization (WHO) and the Declaration of Helsinki 1964 and subsequent amendments or comparable ethical standards.

### Data availability statement

All the data generated or analyzed during this study are included in this published article (and its Supplementary Information files).

## Supplementary Tables

**Table S1:** Differentially altered metabolites in the RRMS group compared to the HS group

**Table S2:** Differentially altered metabolites in the PPMS group compared to the HS group

**Table S3:** Differentially altered metabolites in MS (RRMS and PPMS) compared to HS

